# Generative Models for Prediction of Non-B DNA Structures

**DOI:** 10.1101/2024.03.23.586408

**Authors:** Oleksandr Cherednichenko, Maria Poptsova

## Abstract

**Motivation:** Deep learning methods have been successfully applied to the tasks of predicting non-B DNA structures, however model performance depends on the availability of experimental data for training. Experimental technologies for non-B DNA structure detection are limited to the subsets that are active at the time of an experiment and cannot detect entire functional set of elements. Recently deep generative models demonstrated promising results in data augmentation approach improving classifier performance trained on augmented real and generated data. Here we aimed at testing performance of diffusion models in comparison to other generative models and explore the data augmentation approach for the task of non-B DNA structure prediction.

**Results:** We tested denoising diffusion probabilistic and implicit models (DDPM and DDIM), Wasserstein generative adversarial network (WGAN) and vector quantised variational autoencoder (VQ-VAE) for the task of improving detection of Z-DNA, G-quadruplexes and H-DNA. We showed that data augmentation increased the quality of classifiers with diffusion models being the best for Z-DNA and H-DNA while WGAN worked better for G4s. Diffusion models are the best in diversity for all types of non-B DNA structures, WGAN produced the best novelty for G-quadruplexes and H-DNA. Since diffusion models require substantial resources, we showed that distillation technique can significantly enhance sampling in training diffusion models. When considering three criteria -quality of generated samples, sampling speed, and diversity, we conclude that trade-off is possible between generative diffusion model and other architectures such as WGAN and VQ-VAE.

**Availability:** The code with conducted experiments is freely available at https://github.com/powidla/nonB-DNA-structures-generation.

**Supplementary information:** Supplementary data are available at *Journal Name* online.

## Introduction

The role of non-B DNA structures (quadruplexes (G4), triplexes, Z-DNA, H-DNA, stress-induced duplex destabilization (SIDD), cruciforms), cumulatively called flipons (Herbert et al. 2022), in regulation of cellular processes is currently the subject of extensive studies (Varshney et al. 2020, Wang et al. 2022, Herbert et al. 2023, Umerenkov et al. 2023). The dynamic nature of non-B formation poses limitations on their discovery at the genome-wide level. The existing whole-genome experiments for G4, Z-DNA, H-DNA capture only a subset of all functional flipons in the genome (Shin et al. 2016, Kouzine et al. 2017, Balasubramanian et al. 2018, Marsico et al. 2019, Wu et al. 2020). Deep learning models developed by us and others (Sahakyan et al. 2017, Zhang et al. 2020, Beknazarov et al. 2020, Voytetskiy et al. 2022, Umerenkov et al. 2023) are trained on available experimental datasets and generate whole-genome annotations. The performance of these models is still a subject of improvement.

Recently, generative models like denoising diffusion probabilistic model (DDPM) (Ho et al. 2020), denoising diffusion implicit models (DDIM) (Song et al. 2020), stochastic differential equation based score matching models (SDE) (Yang Song et al. 2020), generative adversarial networks (GAN) (Goodfellow et al. 2014), variational autoencoders (VAE) (Kingma et al. 2013) are increasingly used for data augmentation task to improve deep learning model performance. The goal is to generate new data from a distribution of real data and expand real experimental dataset with artificially generated. Models trained on the augmented dataset presumably have higher performance.

Here we aimed at testing generative approaches on non-B DNA structures -important functional genomic elements for which experimental data is lacking. It is of importance to develop augmentation data approach to improve the quality of existing deep learning models for non-B DNA prediction.

### Formal problem for sequence generation

Here we briefly discuss the problem of generating DNA sequences, borrowing some key ideas from the study (Killoran et al. 2017). A generative model, parameterized by *θ*, tries to learn a real distribution *π*(*x*) and outputs an approximation with distribution *p*(*x*|*θ*) *≈ π*(*x*). The distribution *p*(*x*|*θ*) will be used for sampling. The generator is trained to generate samples similar to real data. So far this task is an unconditional generation task. To generate data with a certain condition, we need to approximate a conditional density *p*(*y*|*x, ?*). Here *y* could be an additional information like properties of a non-B structure or a signal from omics features. There exist various methods for estimation using generative models: likelihood-based with the approximate density (Hoogeboom et al. 2021) realized in diffusion models and variational autoencoders, and free-likelihood with the implicit density (Gulrajani et al. 2017) realized in generative adversarial networks. Another type is likelihood-based with the tractable density realized in autoregressive models and normalizing flows (Bond-Taylor et al. 2021). In this work we consider the first two methods - likelihood-based with the approximate density and free-likelihood with the implicit density.

Different encoding strategies can be used for DNA sequence, consisting of 4 nucleotides A, T, C, G: to work in a discrete space or transform it from a discrete space into continuous. One way is to use one-hot encoding representation that assigns a 4-dimensional vector to each nucleotide, e.g. *A ∼* [1., 0., 0., 0.]^*T*^. In this way a sequence *x* of length *L* is represented as a real-valued matrix of shape (*L*, 4). This approach was used in diffusion models (Wu et al. 2022) and generative adversarial networks (Killoran et al. 2017). Another way is to use a label binarizer and work with a discrete data as it was done in transformer-based Z-DNABERT model (Umerenkov et al. 2023). In this work we decided to use one-hot encoding as it is straight forward to feed into generative pipelines with respect to their initial implementations.

### Generative models for sequence generation

#### Denoising Diffusion Models

Here we briefly describe concepts of utilized generative models. Denoising diffusion models (DDM) are presented in a number of studies (34), (16), (36), (35). We consider denoising diffusion probabilistic model (DDPM) to be a Markov chain model, that gradually removes noise from noised data. Let *q*(*x*) be a forward diffusion process, from where we sample *x*_0_: *x*_0_ *∼ q*(*x*). Let *T* be the number of steps where at each step *t* a small amount of Gaussian noise is added to the sample and *t ∼ 𝒰* [0, 1] where *𝒰* is a uniform distribution. As a result we get a set of *x*_1_, …, *x*_*T*_. Let *β*_1_, …, *β*_*T*_ be a set of parameters to control variance, where each *β*_*t*_ *∈* (0, 1). According to Markov rule the forward process is defined as

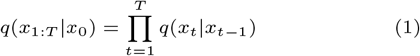

with the Gaussian noise gradually added to the data:

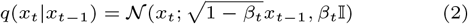

In (16) authors propose to use a reparametrization trick and define the following notations: *α*_*t*_ = 1 *− β*_*t*_ and 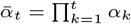, *є ∼ 𝒩* (0, I)

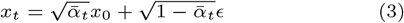

The probabilities for the reverse process:

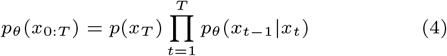

The final transitional probabilities:

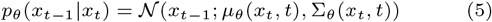

With the following parameters

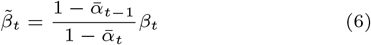

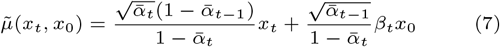

Denoising diffusion implicit model (DDIM) (35) allows to speed up the process by reducing the number of steps. In the framework developed with analog bits (6) the authors present a method of applying Gaussian noise to discrete data encoded via one-hot vectors using transformation to symmetrical real interval [*−*1., 1.]. The objective function is

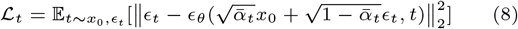

In this work we use framework DNA-Diffusion (8) with the diffusion scheme DDIM. However it is also possible to use DDPM scheme.

#### Wasserstein Generative Adversarial Network

Let’s consider a generative adversarial network concept, starting with defining generator *G*(*z*) and discriminator *D*(*x*). Generator *G*(*z*) takes latent variable *z* and transforms it into a artificial data, then it is applied to discriminator *D*(*x*), which is trained using both real and synthetic data. In the original paper (10) models *D*(*x*) and *G*(*z*) are competing in minmax game by optimizing the objective

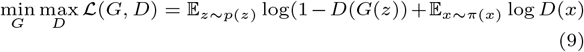

This expression can be reformulated to represent GAN as optimizing Jensen-Shannon divergence, which is symmetric version of KL divergence. In Wasserstein GAN a Wasserstein distance is used

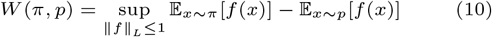

Wasserstain GAN with gradient penalty (WGAN-GP) use gradient penalty instead of clipping (11). The paper proves that a differentiable function is 1-Lipschitz if and only if it has gradients with norm at most 1 everywhere. In WGAN-GP, penalty is used instead of clipping if the gradient norm moves from the target value of 1:

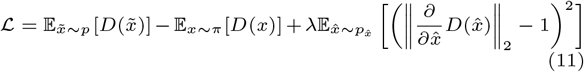

Here 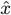 is sampled from 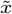 and *x* is sampled from a uniform distribution. WGAN-GP is less prone to mode collapse (37). For that reason we have decided to use this improvement of WGAN since we will use datasets with several modes, i. e. different non-B DNA structures.

#### Vector Quantised-Variational AutoEncoder

Variational Autoencoder (VAE) (22) is a generative model based on the idea of autoencoder rooted in the methods of variational bayesian and graphical model (9). Autoencoder itself is a neural network designed to learn an identity function in an unsupervised way to reconstruct the original input while compressing the data in the process in order to discover a more efficient and compressed representation:

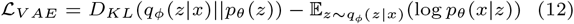

Training process is based on optimizing the following parameters

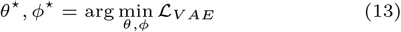

One of the potential improvements of VAE is called Vector Quantised-Variational AutoEncoder (VQ-VAE) (48; 28), and this model learns a discrete latent variable by the encoder, since discrete representations may be a more natural fit for sequence data. Vector quantisation (VQ) is a method to map N-dimensional vectors into a finite set of latent vectors. The process is very much similar to KNN algorithm. The loss function now is represented as

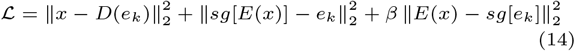

Here outputs *z*_*e*_ and *z*_*q*_ are produced by the encoder and decoder correspondingly, *e*_*k*_ is a vector from a latent embedding space, *sg* is the stop gradient operator (48).

Here we tested three different generative approaches for data augmentation task to improve classifiers for non-B DNA structure prediction. For each method we also explored properties of generated sequences in terms of quality, novelty and diversity.

## Materials and methods

### Data preparation

For Z-DNA we compiled the dataset from experimental Z-DNA ChIP-seq data (Shin et al. 2016, Beknazarov et al. 2020, Voytetskiy et al. 2022), and another experimental Z-DNA dataset obtained with completely different technique named Permanganate/S1 Nuclease Footprinting (Kouzine et al. 2017). For G4 we used experimental ChIP-seq data (Mao et al. 2018). For H-DNA we used experimental dataset generated with Permanganate/S1 Nuclease Footprinting (Kouzine et al. 2017). All DNA sequences containing non-B DNA structures were centered and adjusted to the length of 512 bp. Each fragment was labeled as containing a non-B DNA structure (Z-DNA, G4, or H-DNA). For negative labels we took randomly selected regions not overlapping with non-B DNA. We also composed a combined dataset consisted of all three types of non-B DNA structures - Z-DNA, G4s and H-DNA. We used three various methods of classifier validation based on different ways to compose positive and negative classes. First, we feed real data to the detector with randomly sampled negative class. Secondly, we feed artificial data to the detector with shuffled negative class. Thirdly, we feed combined real and artificial data to the detector with shuffled negative labels. We report the average metric obtained with these three methods.

### Model implementation

General schema of data augmentation pipeline with three architectures of generative models and their application to the task of non-B DNA prediction is given in Figure 1.

**Fig. 1.**
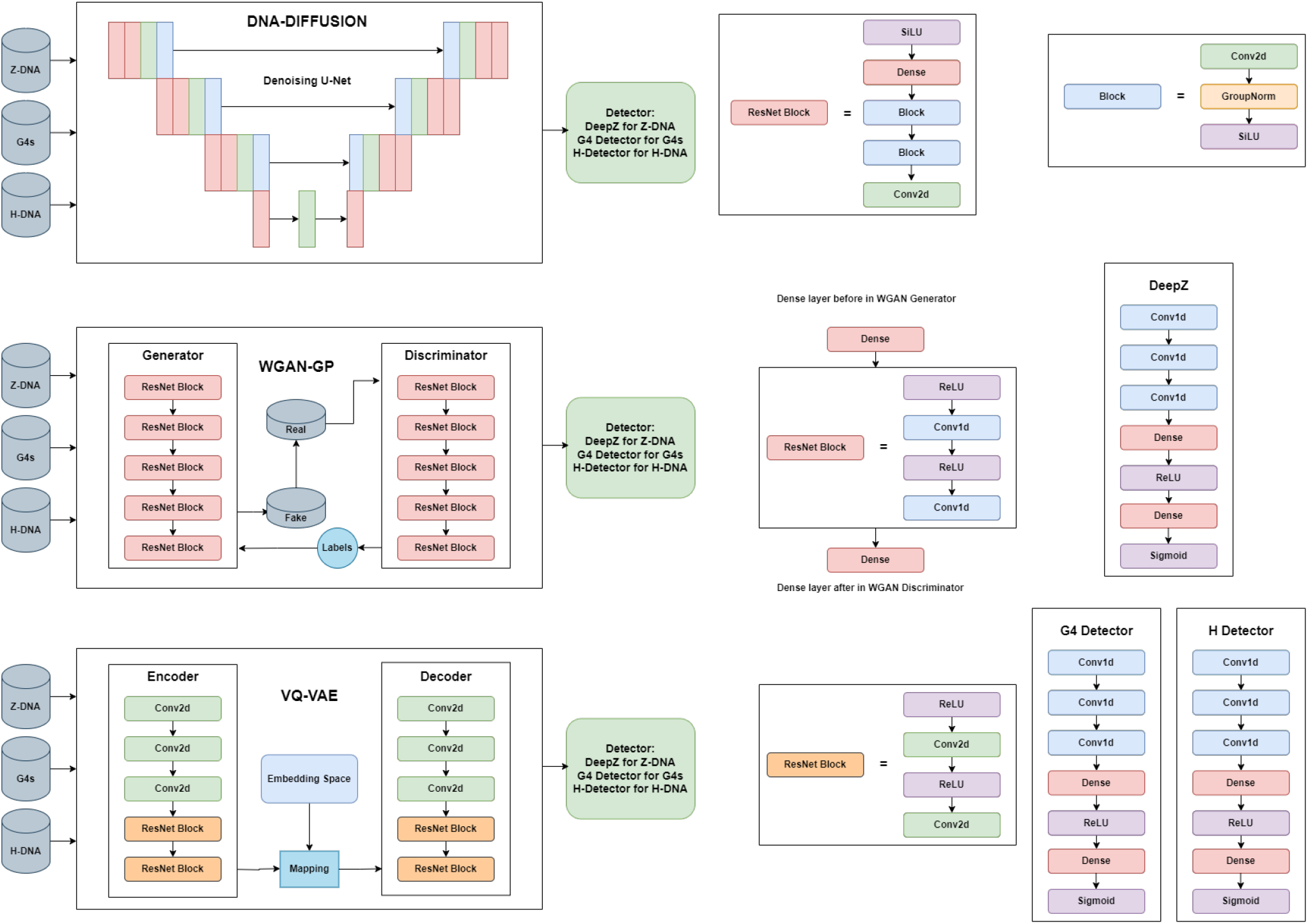
Data augmentation pipeline. Synthetic data are created with three types of generative models: diffusion model, WGAN, and VQ-VAE. Concatenated real and artificial data are fed to the detector for testing the power to identify non-B DNA structures.

#### Diffusion model

The U-net-based diffusion model consists of 4 ResNet blocks and the block with an initialization convolutional layer with a 7x7 kernel, stride 1 and padding 3, followed by a multi-layer perceptron incorporating learned sinusoidal positional embeddings and linear transformations with GeLU activation. The model includes an embedding layer for labels and employs a series of downsampling and upsampling blocks, each containing ResNet block. ResNet block consists of two blocks with skip connections followed by linear attention mechanisms. The downsampling path involves strided convolutions, while the upsampling path incorporates transposed convolutions. The normalization on convolution blocks is made by GroupNorm layers and SiLU instead of GeLU. This implementation has been originally presented in (DaSilva LF et al. 2024). We have applied minor changes in training loop and validation. The following hyperparameters were used for training: 3 optimizers for diffusion training - AdamW (*α* = 10^*−*4^, *β*_1_ = 0.99, *β*_2_ = 0.99), SGD (*α* = 10^*−*4^, momentum=0.995) and SophiaG(*α* = 10^*−*4^, *β*_1_ = 0.965, *β*_2_ = 0.999, *ρ* = 0.01, weight decay=10^*−*2^), gradient accumulation size = 8. The most stable training was observed under AdamW optimizer. For the loss we considered three options: l1, l2 and huber loss function. Last option allowed us to get faster convergence of losses. The loss function over training epochs and comparison of different loss functions for diffusion model are presented in Supplementary Figures 1-3.

#### WGAN

WGAN architecture is composed of a generator and a discriminator, and each of these networks consists of five ResNet blocks. Each block consists of a convolutional layer with 5x5 kernel and padding 2, followed by ReLU activation. For the training of WGAN we used the following hyperparameters: loss with gradient penalty *λ* = 10, model size is 32, trained over 100 epochs via AdamW optimizer with *α* = 0.01, *β*_1_ = 0.9, *β*_2_ = 0.99. Training of WGAN model was accompanied by calculation of two edit distances (see Supplementary Figures 4-5). *D*_*self*_ is the average edit distance in a set of 1000 generated sequences where distance is measured between the sequence and its nearest neighbor (except for itself). This metric is also measured in the test and training datasets. If generated data is as diverse as the data in the training and test datasets, the values of *D*_*self*_ in the generated sequences will be similar. *D*_*train*_ is the average distance that is used to measure GAN training. *D*_*train*_ will also be measured in the test dataset. The low value of this metric alerts the overfitting of the GAN on the training set. Along with the edit distances we provide expressions for various costs functions that we used for training. The cost function for training the discriminator aims to minimize the difference between the Wasserstein distance for real and generated samples: cost = *D*_*fake*_ *− D*_*real*_ + gradient penalty. Wasserstein cost for training the discriminator is the difference between the average output of the discriminator for real and generated samples: wasserstein cost = *D*_*real*_ *− D*_*fake*_. The cost function for validating the discriminator is computed similarly to the training cost: cost validation = *D*_*valfake*_ *− D*_*valreal*_. The Wasserstein cost function for validating the discriminator is computed similarly to the training cost: cost validation = *D*_*valreal*_ *− D*_*valfake*_. The cost function for training the generator aims to maximize the average output of the discriminator for generated samples: gen cost = *−D*_*fake*_.

#### VQ-VAE

VQ-VAE architecture consists of encoder and decoder models. Both encoder and decoder consist of 2 strided convolutional layers with stride 2 and window size 4 × 4, followed by two residual 3 × 3 blocks (implemented as ReLU, 3x3 conv, ReLU, 1x1 conv), all having 256 hidden units. The decoder similarly has two residual 3 × 3 blocks, followed by two transposed convolutions with the stride 2 and the window size 4 × 4. Transformation into discrete space is performed by VectorQuantizerEMA with embedding (512, 64). Training is executed with AdamW (*α* = 0.01, *β*_1_ = 0.9, *β*_2_ = 0.99) for 200 epochs, the embedding dim is set to 64, commitment cost is 0.25, decay for moving average is set to 0.999. The training curves are presented in Supplementary Figure 6.

### Model Training

Training of all models was implemented using the combined dataset of non-B structures and separately datasets for Z-DNA, G4s, and H-DNA. We tested several ratios of negative to positive classes. Training and test sets were split in the proportion of 70:30 maintaining the same ratio on different chromosomes. Below we briefly describe model’s structures and training process.

Diffusion architecture core consists of a large U-net model with attention adopted from (DaSilva LF et al. 2024). Diffusion model were trained using approach of exponential moving average as it was suggested in the initial implementation (Chen et al. 2022), an accelerator, which provides an opportunity to use gradient accumulation to address large batch problem given a limited amount of memory, and multi-gpu parallel training.

For optimization of Wasserstein model we used WGAN-GP function. During the training we calculated the score between K-nearest neighbours by evaluating two edit distances. For optimization of training, nearest neighbours were determined with KDTree data structure.

VQ-VAE consists of encoder and decoder models. We used VectorQuantizerEMA as a vector quantization module for the embedding representation. After the embedding representation is obtained, it is fed to the decoder model. In addition to the initial objective loss of VQ-VAE (van den Oord et al. 2017), we used perplexity as another metric of evaluating the reconstructed data as described in (DaSilva LF et al. 2024). Low perplexity indicates that the model is capable of reconstructing an input with respect to similarity. high perplexity means the opposite. With this approach we select the model with optimal values of parameters, which does not suffer from overfitting. Impact of exponential moving average on training stability of diffusion model is given in Supplementary Figure 7.

### Progressive distillation for diffusion model

One of the challenges that we faced while working with diffusion model was its inference speed. It takes about 8 hours to sample 50k sequences via trained diffusion model using DDIM sampling technique. To address this challenge we decided to test the method of progressive distillation of diffusion model (Salimans et al. 2022). For the detailed description of this algorithm we refer to the work (Meng et al. 2022) where authors presented the method how to distillate classifier guided diffusion. We implemented naive version of distillation process and present the results in Supplementary Tables 1-3. We adopted this method by training one student model, and this student model became a teacher only once. The implementation of this algorithm and the loss for the student model were borrowed from (49). First, we train one diffusion model and we call it a teacher model. Secondly, we copy this model and evaluate the trained teacher model. Then, we train the copied model and we call it a student model. We can repeat the process for more students models, but it requires a substantial resources to train more models. By implementing only x1 and x2 distillations we already obtained a significant decrease in time of sampling. By x1 distillation we mean that we trained a single student model from a teacher model. For x2 distillation - a student model becomes a teacher to a new student model. However quality metrics for distilled data decreased. As an example, we present heatmaps of such sampled data and its comparison to the original diffusion model (see Figures 2-4).

**Fig. 2.**
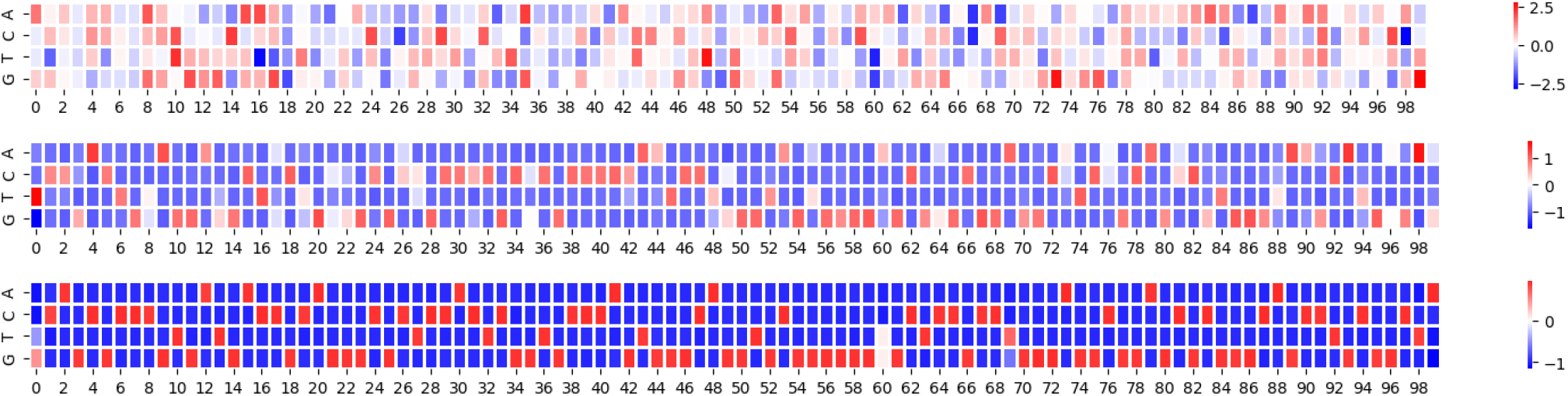
Heatmaps of synthetic sequences, generated by the initial teacher model - a trained diffusion model. First heatmap shows noisy sequences obtained after one step *T* = 1. Second heatmap is a sequence obtained after one hundred steps *T* = 100. Third heatmap is a denoised sequence obtained after one thousand steps *T* = 1000. See explanations in the text.

We also tested a possibility of utilizing less timesteps in inference process. Thus, it should be faster to sample artificial data even having a decrease in quality because of additional noise. We decided this approach worth testing since we also had a slight decrease in quality in the distillation approach. Reducing timesteps allowed us to decrease the sampling time significantly (Supplementary Tables 1-3). For this experiment we used dataset (Mao et al. 2018) with 10k sequences. We used Colab A100 with approximately 90 GB of RAM for the testing time (Supplementary Tables 3-4).

### Evaluation and benchmarks

After deep generative models are trained, we apply them to generate artificial samples of sequences. To test the performance we used the following discriminative models: DeepZ (Beknazarov et al. 2020) for detecting Z-DNA, G4 Detector (Bedrat et al. 2016) to detect G4s, and H Detector with G4 Detector-like architecture to detect H-DNA. We train and evaluate classifier models with the augmented datasets (real plus artificial samples) to estimate performance of generative models. We used different ways to prepare the negative class: random sequences, randomly shuffled sequences from positive class in order to maintain the nucleotide frequency, and by picking sequences according to the pattern (e.g. regular expression), for testing the ability to identify non-B DNA structures.

To evaluate performance of DL models for non-B structure prediction we used conventional metrics for classifiers: ROC-AUC, F1, precision and recall. In addition, we estimated the degree of diversity and novelty of artificial sequences within one dataset and in comparison to the real data. For that we utilized approach presented in (Jain et al. 2022) and calculated novelty and diversity given by the equations (1) and (2) correspondingly. We also provided metrics of performance of DeepZ claimed by the authors in (Umerenkov et al. 2023) using experimental Z-DNA datasets as benchmarks (see Supplementary Table 4).

Formally, the problem of evaluation of generated sequences can be set as follows. Let *𝒟*_0_ be the initial dataset *𝒟*_0_ = *{*(*x*_0_, *y*_0_), …, (*x*_*n*_, *y*_*n*_)*}*, where *x*_*k*_ is a concatenated matrix of encoded sequence, *y*_*k*_ - encoded label for non-B structure. Let *{𝒟*_*j*_ *}* be generated datasets of the same length where j stands for diffusion, WGAN, or VQ-VAE generated sequences. For unconditional generation these datasets can be described as follows: *{𝒟*_*j*_ *}* = *{*(*x*_0_, …, *x*_*n*_)*}*. For conditional generation we expect the output fits the shape of the initial dataset *𝒟*_*j*_ = *{*(*x*_0_, *y*_0_), …, (*x*_*n*_, *y*_*n*_)*}*. We used generated datasets *{𝒟*_*j*_ *}* for augmentation of the initial dataset *𝒟*_0_ in a way of union *𝒟*_*j*_ *∪ 𝒟*_0_ and the labels for generated sequences are assigned by the trained classifiers. We will calculate the following metrics for generated data represented as *{*(*x*_0_, …, *x*_*n*_)*}*, following the same terminology introduced in (Jain et al. 2022)

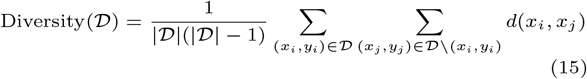

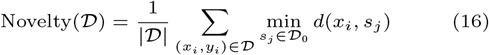

In the original work the distance *d* is defined over a space of discrete objects, so for our purposes we use edit distance as Levenshtein distance between the sequences to evaluate the measure of diversity and novelty. The authors used these metrics inside the training process to evaluate the generated samples. We decided to use these metrics as an additional checkpoint to ensure that an artificial sample satisfies the criteria. As we deal with a huge number of sequences the implementation of these two metrics appeared to be quite challenging. The problem is in the calculation of edit distance, when the loop implementation could be computationally inefficient, and an approximate complexity can be roughly estimated as *O*(*N* ^2^). For efficiency we used KDTree data structure that could be parallelized and reduce the complexity to *O*(log(*N*)).

## Results

### Benchmark of generative models

We tested three generative models - diffusion model (DDIM realization), WGAN and VQ-VAE separately on each type of non-B DNA structure - Z-DNA, G4s and H-DNA, and on the combined dataset (Table 1). The tested datasets for generative models are composed from the combined real and synthetic data, and the results of classification are compared to the classification of real data without augmentation.

**Table 1.**
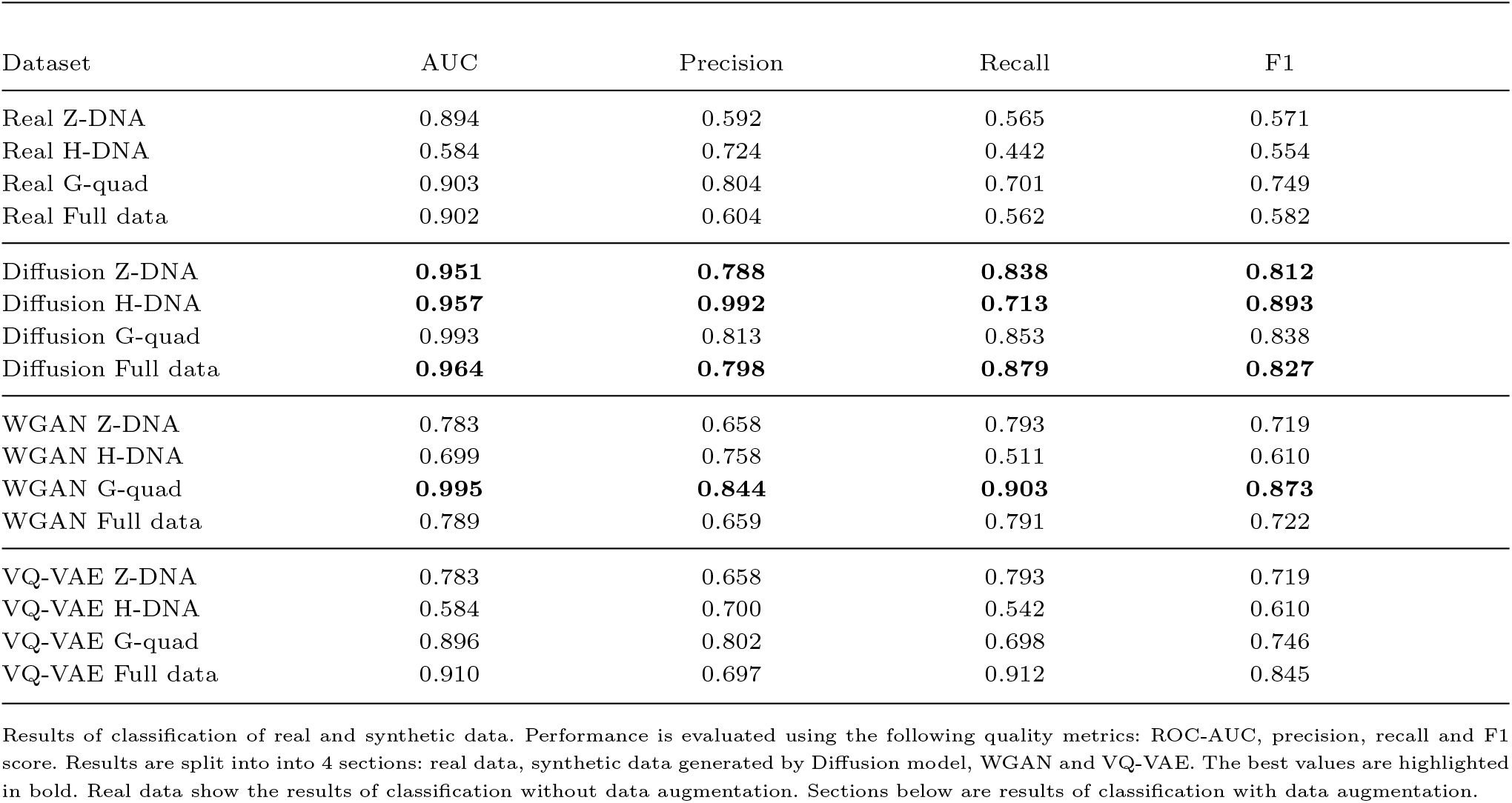
Benchmark of generative models.

| Dataset | AUC | Precision | Recall | F1 |
| --- | --- | --- | --- | --- |
| Real Z-DNA | 0.894 | 0.592 | 0.565 | 0.571 |
| Real H-DNA | 0.584 | 0.724 | 0.442 | 0.554 |
| Real G-quad | 0.903 | 0.804 | 0.701 | 0.749 |
| Real Full data | 0.902 | 0.604 | 0.562 | 0.582 |
| Diffusion Z-DNA | **0.951** | **0.788** | **0.838** | **0.812** |
| Diffusion H-DNA | **0.957** | **0.992** | **0.713** | **0.893** |
| Diffusion G-quad | 0.993 | 0.813 | 0.853 | 0.838 |
| Diffusion Full data | **0.964** | **0.798** | **0.879** | **0.827** |
| WGAN Z-DNA | 0.783 | 0.658 | 0.793 | 0.719 |
| WGAN H-DNA | 0.699 | 0.758 | 0.511 | 0.610 |
| WGAN G-quad | **0.995** | **0.844** | **0.903** | **0.873** |
| WGAN Full data | 0.789 | 0.659 | 0.791 | 0.722 |
| VQ-VAE Z-DNA | 0.783 | 0.658 | 0.793 | 0.719 |
| VQ-VAE H-DNA | 0.584 | 0.700 | 0.542 | 0.610 |
| VQ-VAE G-quad | 0.896 | 0.802 | 0.698 | 0.746 |
| VQ-VAE Full data | 0.910 | 0.697 | 0.912 | 0.845 |

Results of classification of real and synthetic data. Performance is evaluated using the following quality metrics: ROC-AUC, precision, recall and F1 score. Results are split into into 4 sections: real data, synthetic data generated by Diffusion model, WGAN and VQ-VAE. The best values are highlighted in bold. Real data show the results of classification without data augmentation. Sections below are results of classification with data augmentation.

We can see that data augmentation increased the performance of classifiers. Among tested models, the diffusion model has the highest quality metrics. This means that diffusion model produces most accurate synthetic data for Z-DNA, H-DNA and the combined dataset. However, for G4s the best performance was achieved with WGAN model.

### Novelty and diversity of generated non-B DNA structures

We evaluated three generative models for diversity and novelty of generated data (Table 2). Metrics of novelty and diversity are given in (15) and (16) (see Methods). Of all tested models, diffusion model shows the best diversity scores for all type of non-B DNA structures except for the combined dataset. As for novelty, WGAN generates more novel data for G4s and H-DNA. Overall, both metrics confirm that the trained generators are capable of producing novel and diverse data for all types of non-B DNA structures.

**Table 2.** Diversity and novelty for generated data.

| Dataset | Diversity | Novelty |
| --- | --- | --- |
| Real Z-DNA | 73.19 | 0.00 |
| Real H-DNA | 73.19 | 0.00 |
| Real G-quad | 76.96 | 0.00 |
| Real Full data | 74.11 | 0.00 |
| Diffusion Z-DNA | **70.16** | **12.22** |
| Diffusion H-DNA | **76.30** | 9.98 |
| Diffusion G-quad | **74.19** | 11.38 |
| Diffusion Full data | 71.72 | **12.92** |
| WGAN Z-DNA | 69.57 | 11.21 |
| WGAN H-DNA | 69.57 | **10.05** |
| WGAN G-quad | 72.22 | **12.11** |
| WGAN Full data | **72.19** | 11.63 |
| VQ-VAE Z-DNA | 69.51 | 11.62 |
| VQ-VAE H-DNA | 69.99 | 9.98 |
| VQ-VAE G-quad | 69.57 | 11.18 |
| VQ-VAE Full data | 69.31 | 11.80 |

Values in bold indicate the best performance for each type of data.

Independently it is possible to explore diversity of WGAN model by calculating distances (see Methods). The results of final WGAN self and train distances are presented in Table 3 and WGAN train and test distances are presented in Supplementary Figure 4. As we can see, the final absolute values of distances are much higher than 0 at the starting point, and it can be concluded that generated data is diverse.

**Table 3.**
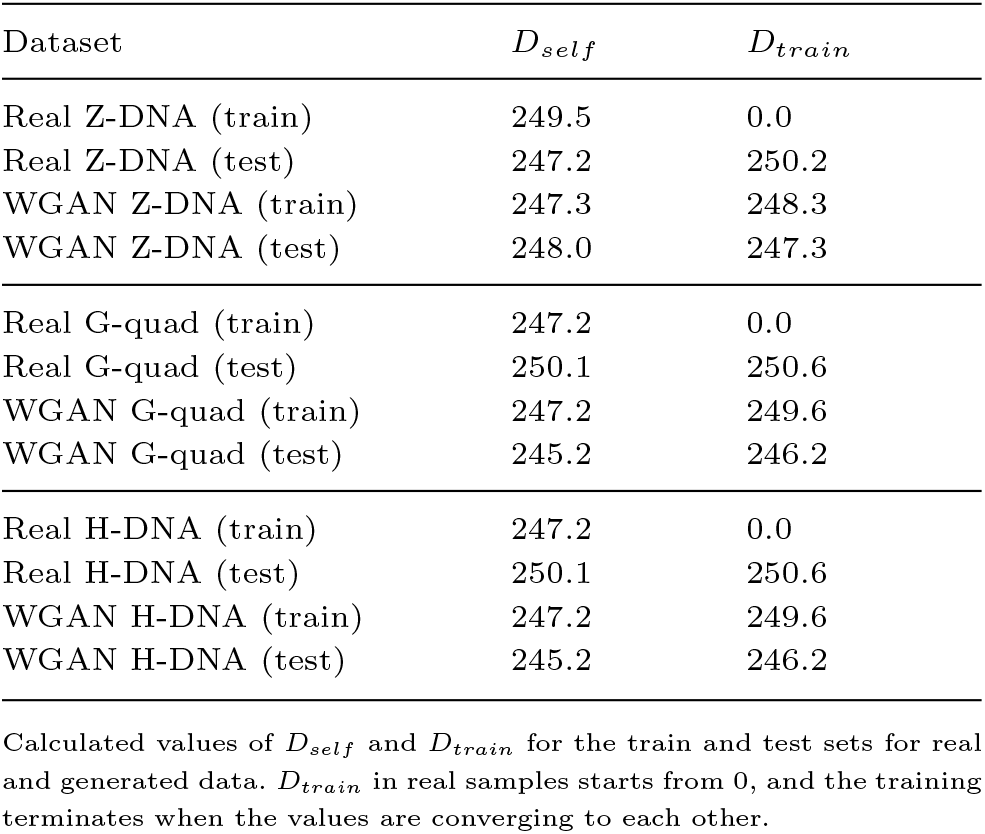
WGAN train and test distances.

| Dataset | $D_{self}$ | $D_{train}$ |
| --- | --- | --- |
| Real Z-DNA (train) | 249.5 | 0.0 |
| Real Z-DNA (test) | 247.2 | 250.2 |
| WGAN Z-DNA (train) | 247.3 | 248.3 |
| WGAN Z-DNA (test) | 248.0 | 247.3 |
| Real G-quad (train) | 247.2 | 0.0 |
| Real G-quad (test) | 250.1 | 250.6 |
| WGAN G-quad (train) | 247.2 | 249.6 |
| WGAN G-quad (test) | 245.2 | 246.2 |
| Real H-DNA (train) | 247.2 | 0.0 |
| Real H-DNA (test) | 250.1 | 250.6 |
| WGAN H-DNA (train) | 247.2 | 249.6 |
| WGAN H-DNA (test) | 245.2 | 246.2 |

Calculated values of $D_{self}$ and $D_{train}$ for the train and test sets for real and generated data. $D_{train}$ in real samples starts from 0, and the training terminates when the values are converging to each other.

### Quality of generated non-B DNA structures

We compared properties of generated samples versus real samples. G4s can be classified by loop length that are captured by different expression patterns: regular pattern *G*_3+_*N*_1*−*7_*G*_3+_*N*_1*−*7_*G*_3+_*N*_1*−*7_*G*_3+_and extended pattern *G*_3+_*N*_1*−*12_*G*_3+_*N*_1*−*12_*G*_3+_*N*_1*−*12_*G*_3+_. We calculated G4 regular and extended patterns within the generated and real data and the results are presented in Table 4.

**Table 4.**
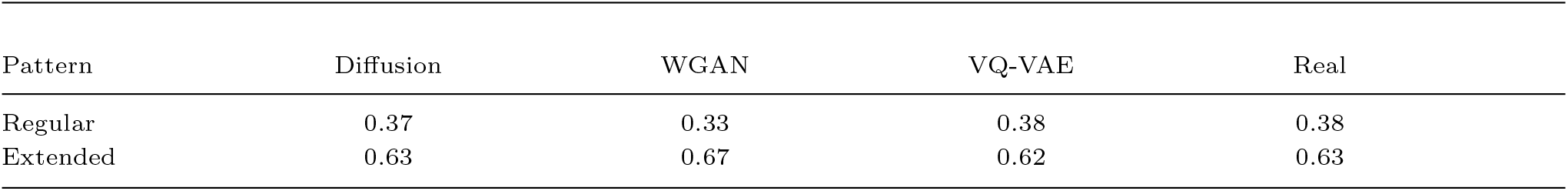
Percent of G4s detected by patterns in generated and real data.

| Pattern | Diffusion | WGAN | VQ-VAE | Real |
| --- | --- | --- | --- | --- |
| Regular | 0.37 | 0.33 | 0.38 | 0.38 |
| Extended | 0.63 | 0.67 | 0.62 | 0.63 |

Percentage-wise all the models behave similar and contain similar proportions of G4s detected by regular or extended patterns. The number of detected patterns in all generative models is very close to the number of detected patterns in real G4 ChIP-seq data with diffusion model being more close to the real data.

One of Z-DNA characteristic property is an alteration of purine-pyrimidine nucleotides that are more prone to switch from B-to Z-conformation. However latest deep learning models based on transformers helped to reveal more complex properties of Z-DNA sequences by analyzing attention scores (Umerenkov et al. 2023). Frequent repeat patterns that are prone to form Z-DNA with corresponding Z-DNABERT attention scores are given in Table 5 together with the occurrence of these repeats in sequences generated by different models. Again we can see that diffusion models generate more close to real data patterns of Z-prone repeats compared to two other generative models.

**Table 5.**
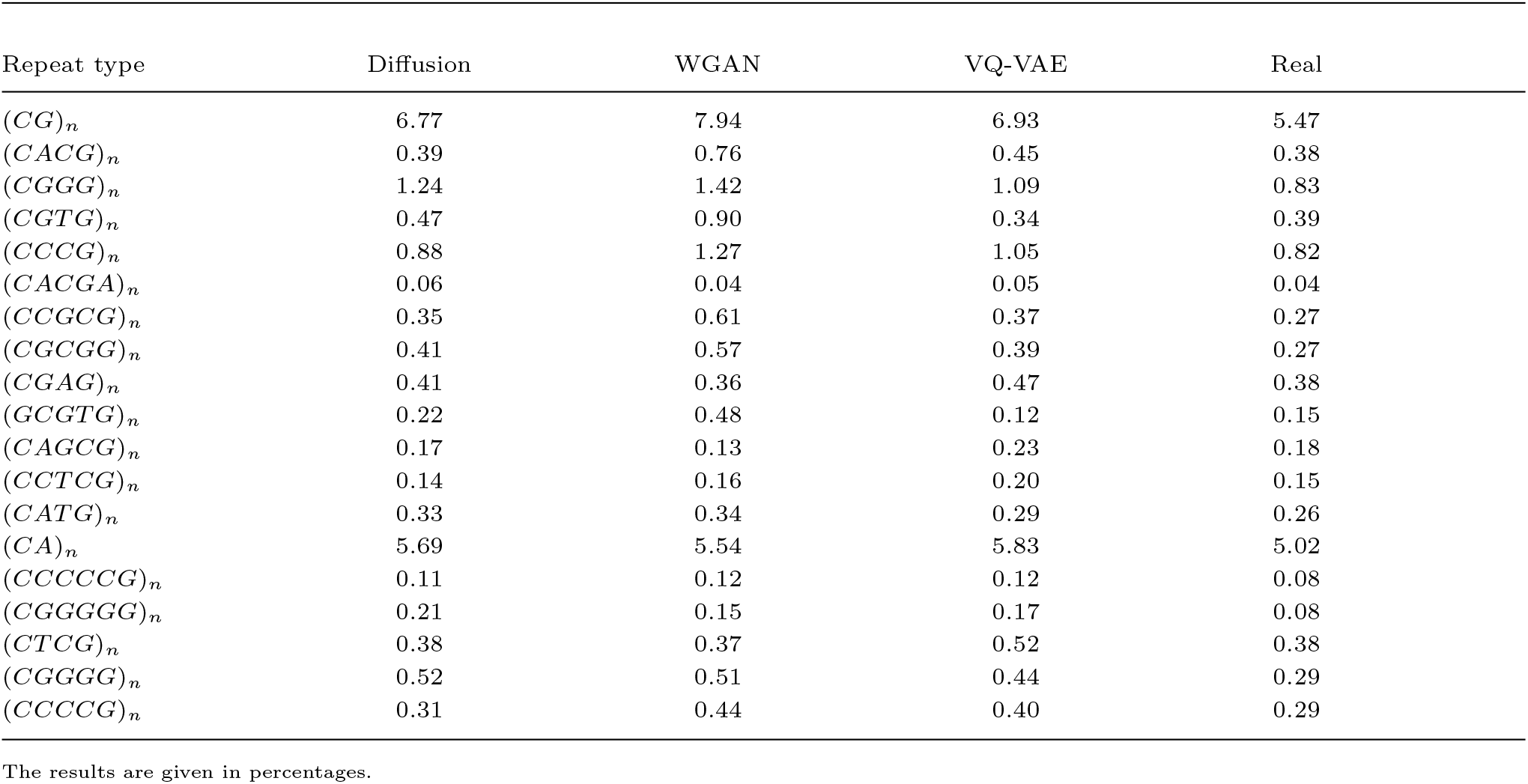
Repeats with Z-DNABERT highest attention scores in generated and real data.

For the evaluation of H-DNA we used Inverted Repeats Finder tool (42) as it was used in (Kouzine et al. 2017) to detect H-DNA. We present results on alignment score in Table 6 with comparison of real and synthetic data.

**Table 6.** Alignment score for inverted repeats in H-DNA in generated and real data.

| Alignment score | Diffusion | WGAN | VQ-VAE | Real |
| --- | --- | --- | --- | --- |
| Mean ± std | $18.1 \pm 3.6$ | $16.9 \pm 1.7$ | $16.7 \pm 1.3$ | $17.9 \pm 4.6$ |

Alignment score produced as an output from Inverted Repeat Finder (IRF)(Warburton et al. 2004). IRF was run with exactly the same parameters as in (Kouzine et al. 2017): minimum alignment scores of 16, matching weight 2, mismatching penalty 6, indel penalty 20, match probability 80, indel probability 10, maximum stem length 100, maximum loop length 8.

### Distillation for diffusion model

For the experiments with distillation we used only G4s dataset since the full dataset is too large, and G4s have strong patterns that can be used for validation of generated data. Here we address an important challenge of low sampling rates in diffusion via simple implementation of distillation techniques (Salimans et al. 2022), (Meng et al. 2022).

The denoising process of diffusion model is presented in Figures 2-4. The figures show heatmaps of a random sample, which is a matrix of 100x4 (sequence length by four nucleotides) where each value represents the measure of certainty for a nucleotide being in a particular position. In the top heatmap one could see the wide range of values, e.g. from -25 to 25 with equal values present in the same position of the sequence. This is a noisy data. After several steps of denoising the range is narrowed to the interval from -1 to 1, as it is required.

Heatmaps of synthetic sequences, generated by the initial teacher model, which is a trained diffusion model, are presented in Figure 2. First heatmap shows noisy sequences obtained after one step *T* = 1 with a high range of values. In this case, the variance from initial encoding is of the same order of magnitude. Second heatmap is a sequence obtained after one hundred steps *T* = 100, and it is more close to the initial range of values. Third heatmap is a denoised sequence obtained after one thousand steps *T* = 1000, and it represents a synthetic sample.

Figure 3 presents heatmaps of synthetic sequences, generated by the first student model - a diffusion model that was trained with the evaluation of the teacher model. First heatmap shows noisy sequences obtained after one step *T* = 1 with high variance from the initial range of transformation that was applied to the sequences, i.e. rescaling to [*−*1., 1.]. We can see that after one step of denoising we obtained a range much higher than for the teacher model. This result is expected since the model was trained on the outputs that were obtained after more timesteps. Second heatmap is a denoised sequence obtained after *T* = 100. It is more close to the initial range of values. Third heatmap is the denoised sequence obtained after one thousand steps *T* = 1000. We can see that the quality is sufficient and the student model generates good samples.

**Fig. 3.**
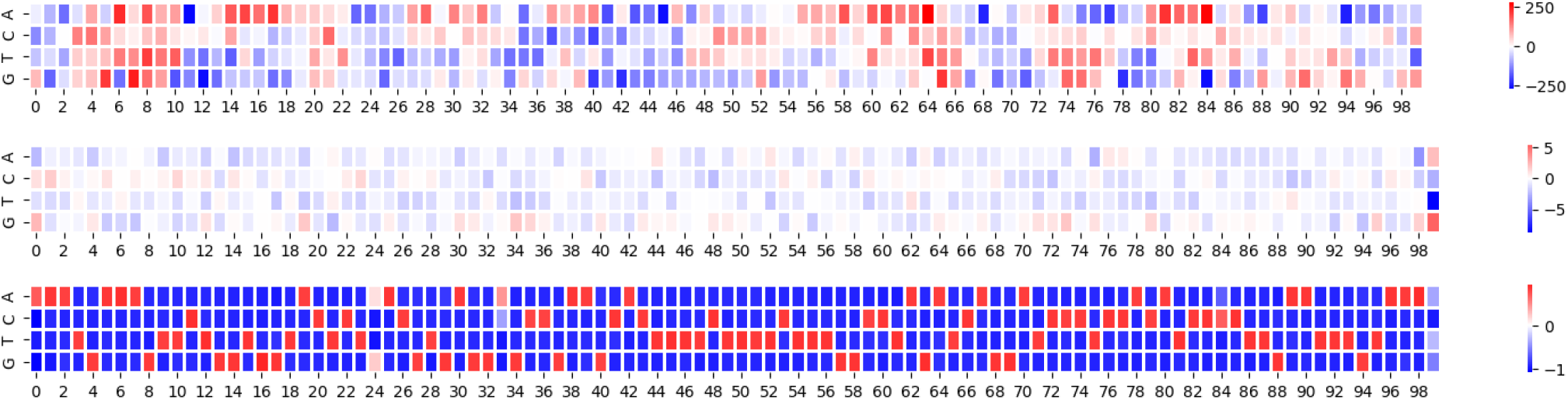
Heatmaps of synthetic sequences, generated by the first student model - a diffusion model that was trained with the evaluation of the teacher model. First heatmap shows noisy sequences obtained after one step *T* = 1. Second heatmap is a denoised sequence obtained after *T* = 100. Third heatmap is the denoised sequence obtained after one thousand steps *T* = 1000. See explanations in the text.

Figure 4 presents heatmaps of synthetic sequences, generated by the second student model - a diffusion model that was trained with evaluation of the first student model in the role of the teacher model. First heatmap shows a noisy sequence obtained after one step *T* = 1 with a variance lower than that of the first heatmap of the first student model (Figure 3), e.g. values are close to interval [*−*20, 20] (compared to [*−*250, 250] in Figure 3). Also in the first heatmap we see that only some positions have high level of uncertainty, e.g. high values close to boundaries of the interval [*−*20, 20]. Second heatmap is a sequence obtained after one hundred steps *T* = 100. Third heatmap is a denoised sequence obtained after one thousand steps *T* = 1000. We can see that this heatmap is close to the third heatmap depicted in Figures 2-3. This proves that even 2 steps of distillation could be sufficient for our task.

**Fig. 4.**
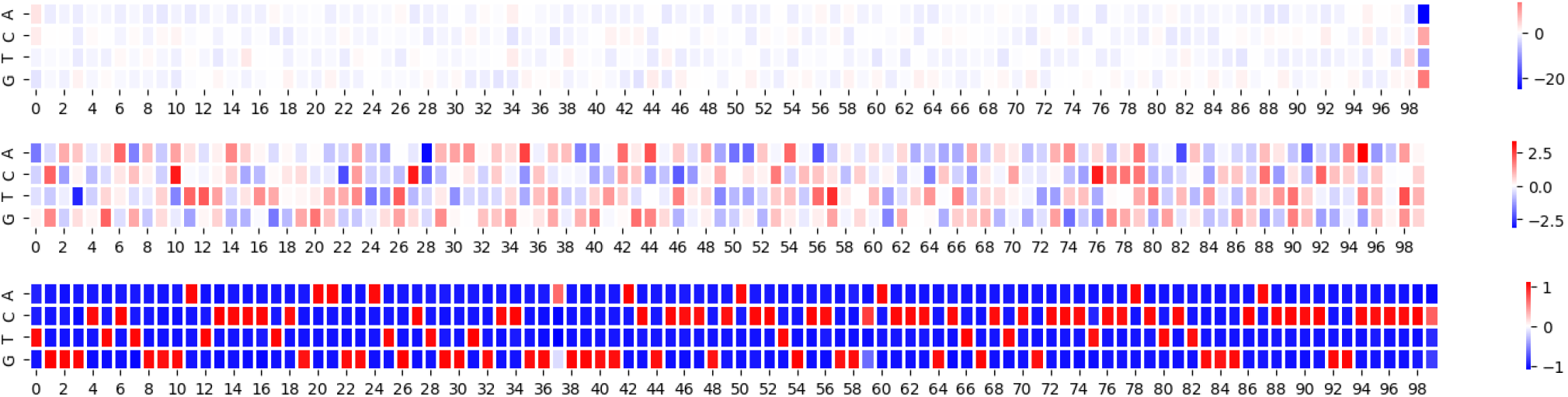
Heatmaps of synthetic sequences, generated by the second student model - a diffusion model that we trained with evaluation of the first student model in a role of the teacher model. First heatmap shows a noisy sequence obtained after one step *T* = 1. Second heatmap is a sequence obtained after one hundred steps *T* = 100. Third heatmap is a denoised sequence obtained after one thousand steps *T* = 1000. See explanations in the text.

The obtained results justify the usage of student models to generate denoised samples. To test how good are synthetic data produced by student models we applied it to discriminator models and evaluated the same quality metrics as in Supplementary Table 2. The results for distilled models are presented in Supplementary Tables 3-4. We conclude that the distillation approach can significantly enhance the sampling process while obtaining fair quality. However, usage of this method with larger datasets requires substantial computational resources because each step in distillation algorithm corresponds to training of a diffusion model.

### Validation of generative approach on mouse data

Finally, we repeated the same set of experiments for mouse genome data (see Supplementary Tables 5-6) and benchmarked the generative model for non-B structures in mouse genome. The results are qualitatively repeated the results presented here for human genome with diffusion model being superior on mouse datasets.

## Discussion

Diffusion models created a new paradigm in generative deep learning. They became SOTA models in various fields such as computer vision (Croitoru et al. 2022), natural language processing (Shin et al. 2023), speech recognition (Zhang et al. 2023). It is still a subject of ongoing research to estimate the performance of diffusion models in generation of functional genomic sequences and compare them with other generative approaches. Since training diffusion models are more computationally efficient compared to other generative approaches it is important to understand which models can be used to replace the diffusion models for a particular task if the quality of generated data is not significantly worse.

In this work we present the application of recently developed framework DNA-Diffusion (DaSilva LF et al. 2024) for the task of generating an important class of functional genomic elements - non-B DNA structures. We benchmarked diffusion models with alternative methods in generative deep learning such as Wasserstein generative adversarial networks and vector quantized variational autoencoder.

Our results showed that diffusion model showed high performance outperforming WGAN and VQ-VAE models on almost all datasets. However, on G4s WGAN model performed better than the diffusion model (see Table 1). The difference between G4s and other non-B structures is that G4s have a characteristic sequence pattern. For that reason a classifier model shows high performance on G4 even without augmentation with ROC-AUC higher than 0.9. Generative adversarial networks are often have a mode collapse problem and this may explain the fact that WGAN model performed high on quadruplexes. Since diffusion model requires more resources WGAN model can be used to generate quadruplexes to maintain a tradeoff between resources and quality.

We evaluated two metrics of diversity and novelty for artificial data. These metrics are based on the edit distance calculation and calculated only for generated sequences to determine how different they are from the real samples. Additionally, these metrics reflect how synthetic sequences vary among each other. We demonstrated that diffusion model showed the best average diversity and novelty among all four datasets. However, for Z-DNA the difference is small, and both WGAN and VQ-VAE show good results. This could be due to the captured pattern of Z-DNA-prone repeats detected by Z-DNABERT attention scores (see Table 5). For H-DNA we observe lower novelty and the synthetic sequences with H-DNA are similar to real and to each other. We assume this is because for H-DNA we used only one source of experimental data (Kouzine et al. 2017), while for Z-DNA we used two orthogonal experimental datasets. G4s also have a strong regular pattern (see Table 4) that ensures that sequences are novel.

Further work in this direction may include the usage of additional data for training deep learning models. For example, it is known that G4s can belong to different classes (Ma et al. 2020) and we can train a conditional generator that will generate G4s of a certain class. The proposed approach can be applied to other types of non-B DNA structures such as triplexes and i-motifs. Generative approach might help to conduct more comprehensive analysis of functional role of these structures and overcome the problem of insufficient experimental data.

While evaluating different generative models we considered the concept of generative trillema (Zhisheng et al. 2021). The concept states that models should be compared based on three criteria: quality of samples, sampling speed, and diversity. In terms of these metrics there is no one superior model architecture. Diffusion models are good with quality and diversity, but low in a sampling speed. WGAN is not high in diversity because of mode collapse effect. VQ-VAE could experience problem with quality in comparison to two other models. This is partially the reason why we used standalone metrics of diversity (15) and novelty (16) to evaluate artificial samples. For the sequences of large length, in comparison to images, it is much harder to evaluate how the samples are different from each other.

In addition, we also tested both DDIM and DDPM sampling techniques. In (Chen et al. 2022) authors show that both are applicable to their framework. Since we are interested in tradeoff between fast sampling and quality, DDIM is more convenient because it allows to produce synthetic data faster. We did not conduct experiments to determine how the choice of the scheme affects the results. We relied on the results obtained by (Chen et al. 2022) that show the high quality of sampling for both methods. However, to address the important challenge of enhancing sampling it was the progressive distillation approach that allowed us significantly reduce inference time.

## Data availability

The code and data underlying this article are available at https://github.com/powidla/nonB-DNA-structures-generation

## Supplementary data

Supplementary data are available.

