## Supplementary Figures and Tables for "Generative Models for Prediction of Non-B DNA Structures"

### Supplementary Materials

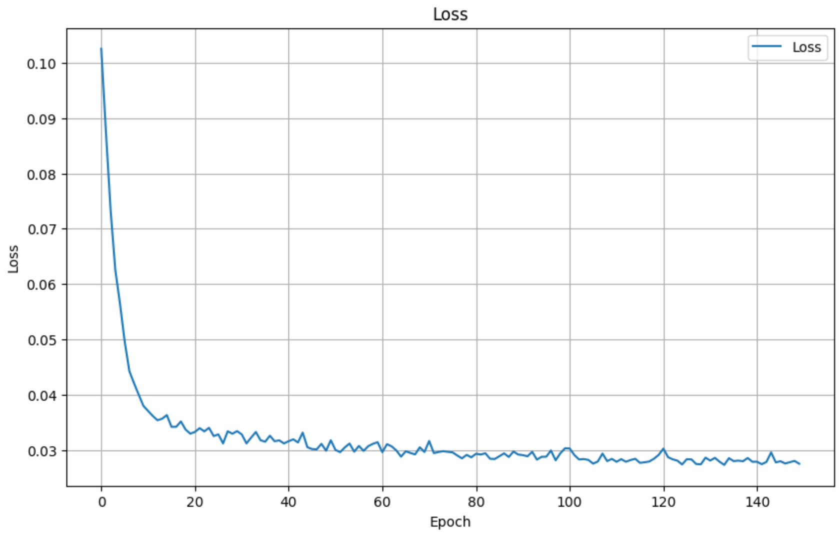


Supplementary Figure 1: Training loss function for diffusion model on full dataset: Z-DNA, G4s, and H-DNA. Here EMA and noising scheduler $\beta(t) = 60t^{17} + 8t^3 + 4t$ work better than linear and cosine schedulers. It makes training more stable.

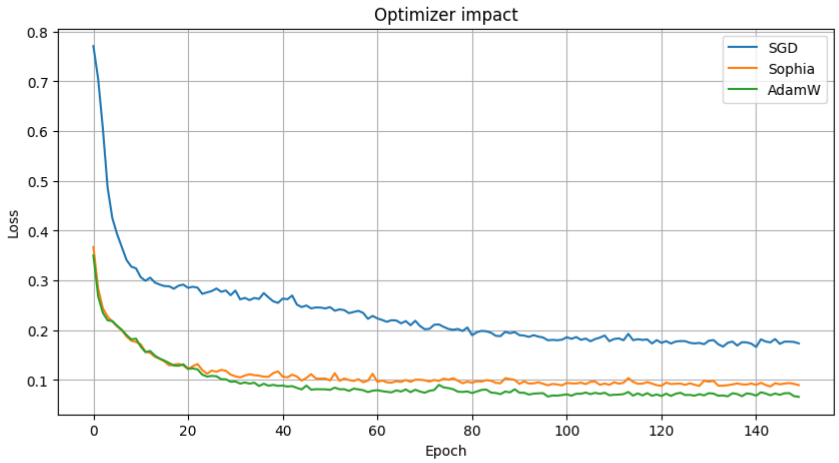


Supplementary Figure 2: Losses convergence with various optimizers and best parameters for each optimizer. Three various optimizers were tested: AdamW ($\alpha = 10^{-4}, \beta_1 = 0.99, \beta_2 = 0.99$), SGD ($\alpha = 10^{-4}$, momentum=0.995) and SophiaG($\alpha = 10^{-4}, \beta_1 = 0.965, \beta_2 = 0.999, \rho = 0.01$, weight decay=$10^{-2}$).

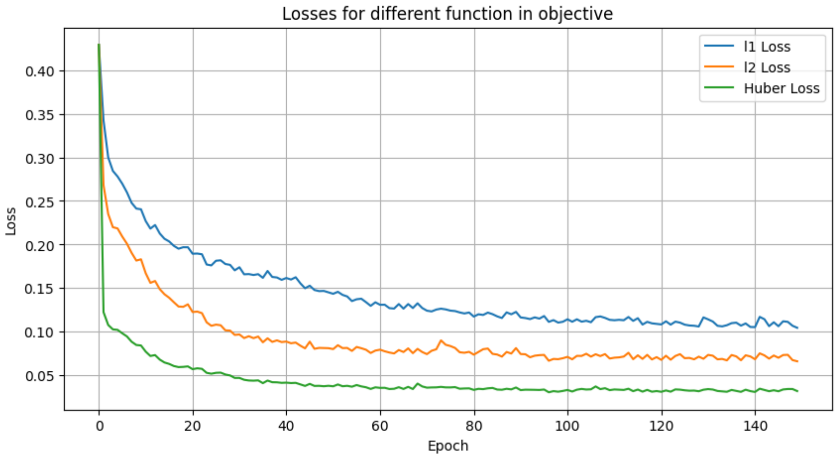


Supplementary Figure 3: Training losses with different functions in objective function for diffusion model. Huber function has faster descent to the average loss value.

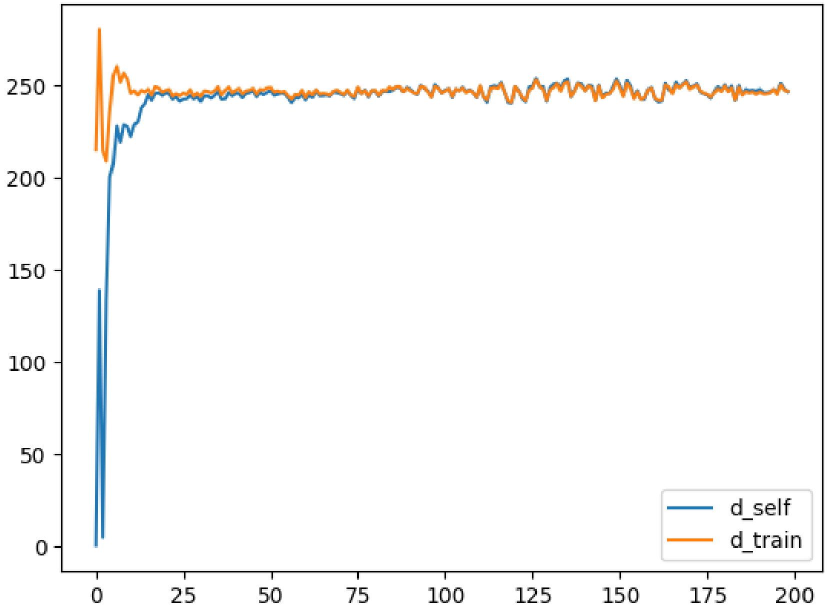


Supplementary Figure 4: Training curves of WGAN model. When the values converge to each other, the model has finished learning patterns. The absolute value of final distance is more than zero, which ensures that the model produces diverse samples.

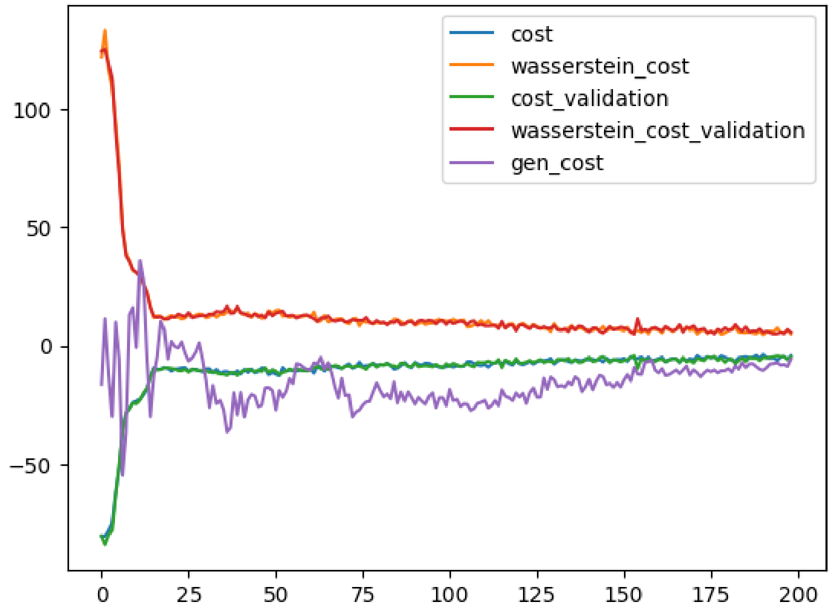


Supplementary Figure 5: Cost functions during the training of WGAN model. All the curves converge to zero over the training process.

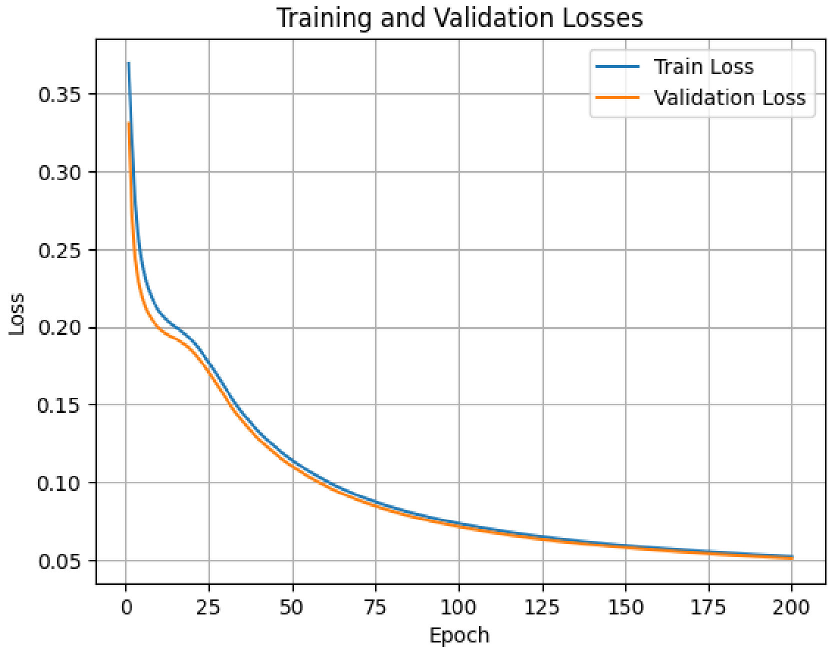


Supplementary Figure 6: Training curves for VQ-VAE model.

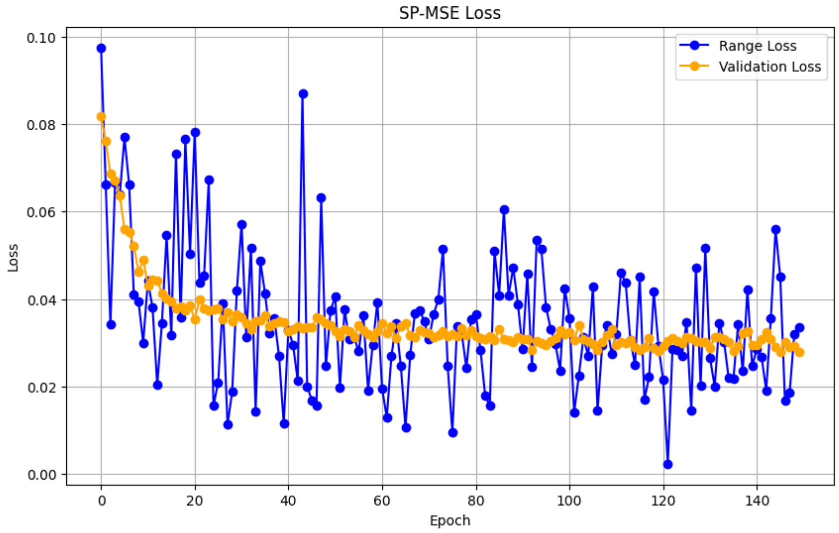


Supplementary Figure 7: Difference between validation curves with and without exponential moving average. We used cosine scheduler to show a possible instability of the training process. Range loss corresponds to the curve without exponential moving average.

Supplementary Table 1: Impact of timesteps $T$ on classifier performance

| Model | AUC | Precision | Recall | F1 |
|---|---|---|---|---|
| not dist | **0.985** | **0.769** | **0.935** | **0.844** |
| dist x2 | 0.872 | 0.804 | 0.712 | 0.755 |
| dist x4 | 0.873 | 0.808 | 0.830 | 0.812 |
| $T = 1$ | 0.388 | 0.556 | 0.421 | 0.479 |
| $T = 10$ | 0.724 | 0.723 | 0.626 | 0.671 |
| $T = 100$ | 0.872 | 0.804 | 0.712 | 0.755 |
| $T = 1000$ | **0.985** | **0.769** | **0.935** | **0.845** |

Performance of classifier on synthetic data generated by models with various timesteps and various distilled models. In this experiment we used only G4s dataset. Results are split into two groups: synthetic data generated by distilled models and synthetic data generated by models with different amount of time steps. Values in bold designate the best metric in each group.

Supplementary Table 2: Diversity and novelty for evaluating impact of timesteps $T$

| $T$ | Diversity | Novelty | Time (min) |
|---|---|---|---|
| 1000 | 76.24 | 11.95 | 119.15 |
| 100 | 75.74 | 11.33 | 37.10 |
| 10 | 72.24 | 11.32 | 4.50 |
| 1 | 65.71 | 9.03 | 0.86 |

Diversity and novelty for various timesteps in sampling, time represents the sampling speed. The experiments were conducted using Colab A100 with approximately 90 GB of RAM. These metrics were calculated for real and generated samples. In this experiment we used only G4s dataset. As expected, for one sampling step we have pure noise, however metrics of novelty and diversity are close to the real samples. The difference in time is significant for fast sampling purposes.

Supplementary Table 3: Diversity and novelty for evaluating distillation impact

| $T$ | Diversity | Novelty | Time (min) |
|---|---|---|---|
| not dist | 76.24 | 11.95 | 119.15 |
| dist x2 | 76.31 | 12.08 | 37.10 |
| dist x4 | 77.04 | 12.04 | 12.98 |

Diversity and novelty metrics for various timesteps in sampling. Time represents the sampling speed. The experiments were conducted using Colab A100 with approximately 90 GB of RAM. These metrics were calculated for real and generated samples. In this experiment we used only G4s dataset. The difference for distilled models in time is significant and it allows using student models for sampling in order to reduce time resources maintaining relatively the same quality.

Supplementary Table 4: DeepZ performance

| Metric | Dataset | Value |
|---|---|---|
| ROC-AUC | Z-DNA Dataset 1 | 0.94 |
| Precision | Z-DNA Dataset 1 | 0.59 |
| Recall | Z-DNA Dataset 1 | 0.56 |
| F1 | Z-DNA Dataset 1 | 0.57 |
| ROC-AUC | Z-DNA Dataset 2 | 0.89 |
| Precision | Z-DNA Dataset 2 | 0.01 |
| Recall | Z-DNA Dataset 2 | 0.30 |
| F1 | Z-DNA Dataset 2 | 0.02 |

DeepZ was used as as a classifier for finding functional regions of Z-DNA in (Beknazarov et al., 2019). Z-DNA Dataset 1 - (Kouzine et al., 2017) and Dataset 2 - (Shin et al., 2016). Values of model's performance were taken from (Beknazarov et al., 2019).

Supplementary Table 5: Diversity and Novelty for generated samples in mouse genome

| Dataset | Diversity | Novelty |
|---|---|---|
| Real Z-DNA | 72.24 | 0.00 |
| Real H-DNA | 64.24 | 0.00 |
| Real Quad | 64.37 | 0.00 |
| Real Full data | 74.11 | 0.00 |
| Diffusion Z-DNA | **73.10** | 11.40 |
| Diffusion H-DNA | 72.24 | 11.95 |
| Diffusion Quad | **74.55** | 12.00 |
| Diffusion Full data | 71.72 | **12.92** |
| WGAN Z-DNA | 72.24 | **12.20** |
| WGAN H-DNA | 72.24 | **12.20** |
| WGAN Quad | 70.21 | **12.88** |
| WGAN Full data | **72.19** | 11.63 |
| VQ-VAE Z-DNA | 71.71 | 11.27 |
| VQ-VAE H-DNA | 73.01 | 11.27 |
| VQ-VAE Quad | 73.00 | 11.27 |
| VQ-VAE Full data | 69.31 | 11.80 |

Values in bold indicate the best performance in each category.

Supplementary Table 6: Benchmark of generative models.

| Dataset | AUC | Precision | Recall | F1 |
|---|---|---|---|---|
| Real Z-DNA | 0.469 | 0.745 | 0.484 | 0.661 |
| Real H-DNA | 0.634 | 0.672 | 0.550 | 0.605 |
| Real Quad | 0.891 | 0.792 | 0.692 | 0.739 |
| Real Full data | 0.902 | 0.604 | 0.562 | 0.582 |
| Diffusion Z-DNA | **0.783** | 0.788 | **0.818** | **0.806** |
| Diffusion H-DNA | 0.918 | 0.806 | 0.776 | 0.790 |
| Diffusion Quad | **0.981** | **0.898** | **0.854** | **0.862** |
| Diffusion Full data | **0.964** | **0.798** | **0.879** | **0.827** |
| WGAN Z-DNA | 0.563 | 0.793 | 0.109 | 0.192 |
| WGAN H-DNA | 0.699 | 0.758 | 0.511 | 0.610 |
| WGAN G-quad | **0.995** | **0.844** | **0.903** | **0.873** |
| WGAN Full data | 0.789 | 0.659 | 0.791 | 0.722 |
| VQ-VAE Z-DNA | 0.549 | **0.899** | 0.142 | 0.245 |
| VQ-VAE H-DNA | 0.834 | 0.817 | 0.713 | 0.761 |
| VQ-VAE Quad | 0.898 | 0.801 | 0.695 | 0.744 |
| VQ-VAE Full data | 0.910 | 0.697 | 0.912 | 0.845 |

Results of classification of real and synthetic data. Performance is evaluated using the following quality metrics: ROC-AUC, precision, recall and F1 score. Results are split into into 4 sections: real data, synthetic data generated by diffusion model, WGAN and VQ-VAE. The best values are highlighted in bold. Real data show the results of classification without data augmentation. Sections below are results of classification with data augmentation.
